# Multi-centered T cell repertoire profiling identifies novel alterations in the immune repertoire of individuals with inflammatory bowel disease and validates previous findings

**DOI:** 10.1101/2025.05.16.654339

**Authors:** Aya K.H. Mahdy, Hesham ElAbd, Érika Endo Kokubun, Valeriia Kriukova, Mitchell Pesesky, Damon May, Christine Olbjørn, Gøri Perminow, May-Bente Bengtson, Petr Ricanek, Svend Andersen, Trond Espen Detlie, Vendel A. Kristensen, Bjørn Moum, Morten Vatn, Jørgen Jahnsen, Bernd Bokemeyer, Johannes Roksund Hov, Jonas Halfvarsson, Stefan Schreiber, the IBSEN and IBSEN-III study group, Bryan Howie, Harlan S. Robins, Marte Lie Høivik, Andre Franke

**Author notes:** Shared first co-authorship (Aya K.H. Mahdy, Hesham ElAbd).

## Abstract

**Introduction:** IBD is an incurable immune-mediated inflammatory disease (IMID), affecting the gut with a high rate of primary- and secondary-loss-of-response to therapy. By investigating the T cell receptor repertoire of individuals with IBD, novel therapeutic and preventive strategies can be identified, and a better understanding of IBD can be obtained.

**Methods:** Whereas most studies have so far focused on the more diverse T cell receptor beta (TRB) repertoire, we here profiled the alpha (TRA) repertoire of three cohorts containing treatment-naïve and treated individuals in addition to individuals living with the disease for >20 years, resulting in an exhaustive dataset containing the TRA repertoire of 2,151 individuals.

**Results:** Using the generated datasets, we were able to replicate previous findings describing the expansion of Crohn’s-associated invariant T (CAIT) cells in individuals with Crohn’s disease (CD) in the three cohorts. Using a hypothesis-free statistical testing framework, we identified clonotypes that were associated with the disease at its different stages, *e.g.,* at the time of diagnosis and decades post-diagnosis. By conducting a meta-analysis across the three cohorts, we were able to identify a set of clonotypes that were associated with the disease regardless of its stage. We validated our findings in a previously published independent test dataset from a German cohort, showing the robustness of the identified sets of clonotypes.

**Conclusion:** The identified clonotypes are potential novel therapeutic targets to treat IBD, *e.g.,* through targeted depletion. These clonotypes are also of major interest as they can be investigated in a targeted fashion to identify culprit antigen(s) in IBD.

## Introduction

Inflammatory bowel disease (IBD) is associated with a significant reduction in the quality-of-life and increased morbidity and is a major burden on the global healthcare system^1,2^. While different therapies have been developed to treat IBD, such as anti-TNF and anti-integrins, they fail to introduce a response in all patients, *i.e.,* primary non-responders^3,4^. Furthermore, loss-of-response is commonly observed as patients develop antibodies against these medications^3^. Thus, there is an urgent need to develop novel therapies that induce long-lasting remission in a large fraction of patients. By depleting T cell clones that are implicated in ankylosing spondylitis (AS), Britanova *et al.*^5^ were able to develop a targeted therapy that introduced remission. Thus, disease-associated T-cell clones are a novel therapeutic target for treating immune-mediated inflammatory diseases (IMIDs) such as IBD.

However, identifying T cell clonotypes that are involved in IBD is not a trivial task for multiple reasons, such as heterogeneities in the clinical presentation, differences in affected tissues, and a complicated genetic architecture, particularly within the human leukocyte antigen (HLA) loci. Multiple HLA alleles have been associated with IBD, for example, HLA-DRB1*03:01^6^, or with one of IBD’s subsets, *i.e.,* Crohn’s disease (CD) and ulcerative colitis (UC), such as HLA-DRB1*07:01, which is strongly associated with CD^7^ and HLA-DRB1*15:01 with UC^8^. Furthermore, each individual experiences a different journey that is characterized by different medication intake as well as different surgeries. By profiling the immune repertoire of individuals with IBD, multiple alterations were identified, such as altered responses toward yeast antigens^9^ and gut microbiota^10^, as well as an expansion of a subset of type II invariant natural killer T cells, particularly in individuals with CD^11^. These cells are termed CAIT cells and have been shown previously to recognize small molecules presented by CD1d proteins^11^. Other alterations have been identified in individuals with IBD, such as the presence of multiple expanded clonotypes in the colonic mucosa of individuals with CD^12,13^. Advances in bulk T cell repertoire sequencing (TCR-Seq)^14^ and statistical analyses have enabled public clonotypes associated with a particular antigenic exposure, such as cytomegalovirus^15^, SARS-CoV2^16^, and Lyme disease^17^, to be identified.

Most of these studies have focused on the more diverse beta chain of the T cell receptor, *i.e.,* the (TRB) repertoire, leaving the less diverse alpha chain repertoire mostly unexplored. From a statistical perspective, studying the TRA repertoire is more promising as a smaller sample size might be needed to identify alterations driving the disease. Furthermore, most unconventional T cells, such as mucosal-associated T (MAIT) cells, are characterized by a semi-invariant TRA chain; thus, the expansion of these cells can be easily quantified from the TRA repertoire. Hence, we aimed here at utilizing TCR-Seq to identify and validate public clonotypes associated with different subsets of IBD at different stages of disease trajectories.

## Material and Methods

### Study cohorts and study design

We profiled the TRA repertoire of three distinct cohorts from Germany and Norway. First, the IBSEN-III cohort (**Table S1**) contains treatment-naive and treated individuals with IBD, in addition to individuals with symptoms of IBD but without any radiological or endoscopic findings, *i.e.,* symptomatic controls (**Fig. 1A**). Second, the IBSEN-20 cohort (**Table S2**), which contains material from patients included in the first IBSEN study^18^, 20 years post-their diagnosis (**Fig. 1A**). The cohort contains 128 individuals with CD and 265 individuals with UC (**Fig. 1A**). From both Norwegian cohorts, PAXgene tubes were collected and used for RNA extraction, subsequently, the RNA was used to profile the TRA repertoire. The third cohort contained 155 individuals with CD and 115 individuals with UC from the German BioCrohn and BioColitis cohorts (BCBC cohort) (**Table S3**), in addition to 199 healthy individuals (**Fig. 1A**). For this German cohort, DNA was extracted from EDTA blood tubes and subsequently utilized to profile the TRA repertoire. After that, we used the hypothesis-free statistical framework described by Emerson and colleagues^19^ to identify sets of clonotypes that are associated either with CD or UC. Subsequently, we performed a meta-analysis on the identified clonotypes from each cohort to identify a robust set of disease-associated clonotypes. Lastly, we used previously published test datasets^11^ to validate the identified CD- and UC-associated clonotypes (**Fig. 1B**).

**Figure 1:**
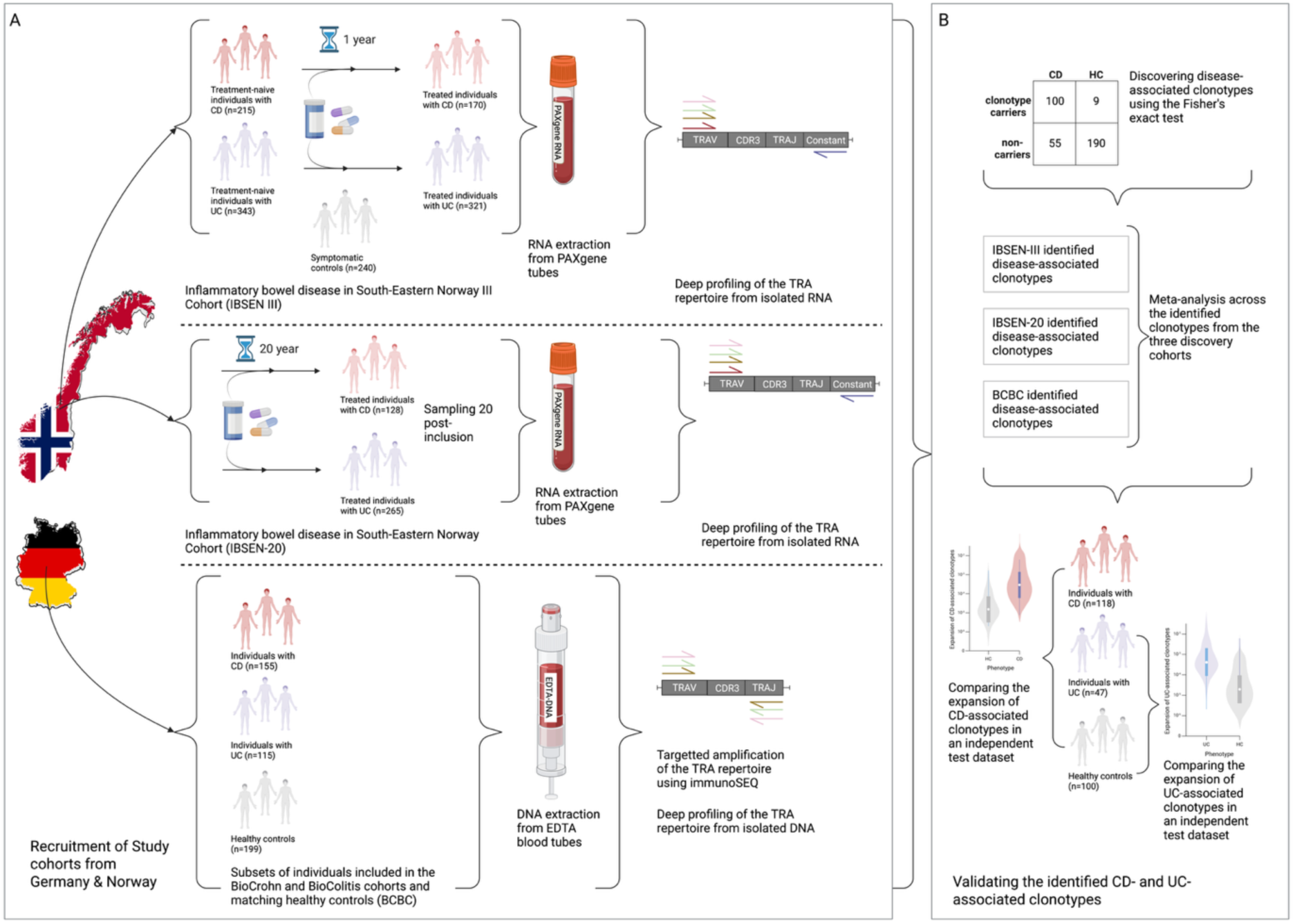
The included cohorts and the analytical pipeline utilized in the current study. (**A**) the pipeline for profiling the TRA repertoire of the three discovery cohorts, namely, IBSEN-III, IBSEN-20, and the BCBC cohorts. (**B**) The analytical framework was used for identifying disease-associated clonotypes from the three discovery cohorts, as well as among the three cohorts using a meta-analysis approach. Lastly, the identified clonotype sets were validated using a previously published test dataset^11^.

### Ethical approvals

The study has been approved by the ethical committee at the University of Kiel under the following ethical votes: D441/16, D474/12, A161/08, A103/14, and A148/14. The IBSEN and IBSEN-III study was approved by the South-Eastern Regional Committee for Medical and Health Research Ethics (Ref: 2010/1540 and 2015/946-3, respectively) and performed in accordance with the Declaration of Helsinki.

### TCR profiling using DNA

The TRA repertoire of the BCBC cohort and the matching healthy controls were profiled using DNA extracted from peripheral blood. Subsequently, up to 18 µg of DNA was used for profiling the TRA repertoire using the immunoSEQ assay (Adaptive Biotechnologies).

### TCR profiling using RNA

The TRA repertoire of the IBSEN-III and the IBSEN-20 cohorts were profiled using RNA extracted from PAXgene tubes collected from peripheral blood. From the IBSEN-III cohort, up to 300 ng of RNA were used, while for the IBSEN-20, up to 200 ng of RNA were used. Subsequently, next-generation sequencing (NGS) libraries of the TRA chain were generated using MiLaboratories’ commercially available kits according to the manufacturer’s instructions. After indexing the samples using Illumina dual indices, samples were pooled together and sequenced using 150 bp paired-read sequencing on the NovaSeq 6000. After demultiplexing the generated sequencing reads per sample, *i.e.,* FASTQ files, they were processed using MiXCR^20^ (V 4.6) to identify and quantify the expansion of the different TRA clonotypes present in each sample.

### Processing the identified clonotypes and generated repertoires

After identifying TRA clonotypes using either immunoSEQ or MiLaboratories kits, we started processing the repertoires by removing non-productive clonotypes, *i.e.,* clonotypes containing a frameshift or a stop codon, and hence they do not encode for a functional TRA chain. Subsequently, we grouped different VJ recombination encodings for the same TRA chain at the protein level into a single clonotype and summed their expansion. That is, each clonotype included in the analysis was a unique VJ recombination with a unique CDR3 amino acid sequence in a sample. Lastly, we removed samples with less than 1000 productive clonotypes.

### Identifying CD and UC- associated clonotypes

To identify TRA clonotypes that are associated with CD or UC, we utilized the framework discussed by Emerson *et al.*^19^, focusing on public clonotypes, *i.e.,* clonotypes present in more than one individual. After identifying public clonotypes, we compared their frequency in cases, *e.g.,* CD or UC, and controls using the one-sided Fisher’s exact test. Subsequently, we used a cutoff of 1x10^-3^ to identify associated clonotypes. For both the IBSEN-III and the BCBC cohorts, we compared the repertoire of CD and UC individuals to symptomatic controls or to healthy controls, respectively, to identify CD and UC-associated clonotypes. Meanwhile, for the IBSEN-20, neither healthy nor symptomatic controls were available, and we compared the repertoire of CD to that of UC and vice versa to identify clonotypes associated with either CD or UC.

### Seeded clustering of TRA-associated clonotypes

To extend the set of disease-associated clonotypes to rarer clonotypes that our study did not have the statistical power to identify, we performed seeded clustering, which involves three main steps.

1. Identifying disease-associated clonotypes using Fisher’s exact test as defined above (**identifying CD and UC-associated clonotypes**). These clonotypes represent the seeds, which are the base for extending the clonotype search on.
2. Extended search: after identifying the seeds, for each seed, we search the entire repertoire for clonotypes that have the same V and J genes as that of the seed and a CDR3 sequence that is at max 1-Levenshtein distance from the seed. The collection of a seed and its similar sequences is referred to as an unpurified meta-clonotype.
3. Seed purification, to purify and prepare the final set of meta-clonotypes, we iterated over each member of the unpurified meta-clonotypes, where we compared the association P-value of the seed and member to that of the seed using Fisher’s exact test. If the P-value of the member and the seed was larger than the P-value of the seed, this member was excluded from the unpurified meta-clonotype. Otherwise, it was kept. The set of clonotypes that survive the purification step, and their seed are referred to as the purified meta-clonotype.

### Performing meta-analysis across the different cohorts

To perform a meta-analysis across the three cohorts described in the study, namely, the IBSEN-III, the IBSEN-20, and the BCBC cohort, we utilized Fisher’s combined P-value approach. After identifying the clonotypes associated with either CD or UC from each cohort independently, *i.e.,* seeds. We focused on the clonotypes that are present in the three cohorts and subsequently calculated an association P-value for each cohort using the one-sided Fisher’s exact test. Thus, we ended up with three P-values for each clonotype that was detected in the three cohorts. These P-values were combined using Fisher’s approach to generate a single P-value. Lastly, we utilized the Benjamini-Hochberg correction method to correct for multiple hypotheses and adjust the P-value. TRA-clonotypes with an adjusted P-value <0.05 were identified as CD- or UC-associated clonotypes identified from the meta-analysis.

### Graph analysis of the identified clonotypes

To perform a network analysis of the identified CD- and UC- associated clonotypes, we used a graph-based approach in which clonotypes were represented as nodes and edges as a similarity metric between these nodes. Two nodes, *i.e.,* TRA clonotypes, were connected by an edge if they shared the same V and J genes and have a hamming distance between their CDR3 amino acid sequences <1. Visualization of the generated graph was performed using Cytoscape^21^.

## Results

### CAIT cells are expanded across different disease stages and are not induced by treatment

We first aimed to study the expansion pattern of Crohn’s-associated invariant T (CAIT) cells in individuals with IBD as well as in healthy controls. Given that CAIT cells can recognize small molecules that resemble drugs and/or bacterial metabolites^22^, we compared the expansion of CAIT cells in treatment-naive individuals with IBD and treated individuals. Using the TRA repertoire of the IBSEN-III cohort, we observed that CAIT cells were expanded in treatment-naive individuals with CD relative to treatment-naive individuals with UC and symptomatic controls (**Fig. 2A**). As the IBSEN-III cohort contains both treatment-naive adults and pediatrics with IBD, these were studies separately. CAIT cells were significantly expanded in adults with CD relative to UC or symptomatic controls (**Fig. 2B**) but not in pediatric cases (**Fig. 2C**). This might be a consequence of differences in the sample size or a true biological difference in the pathogenesis of adult and pediatric forms of IBD.

**Figure 2:**
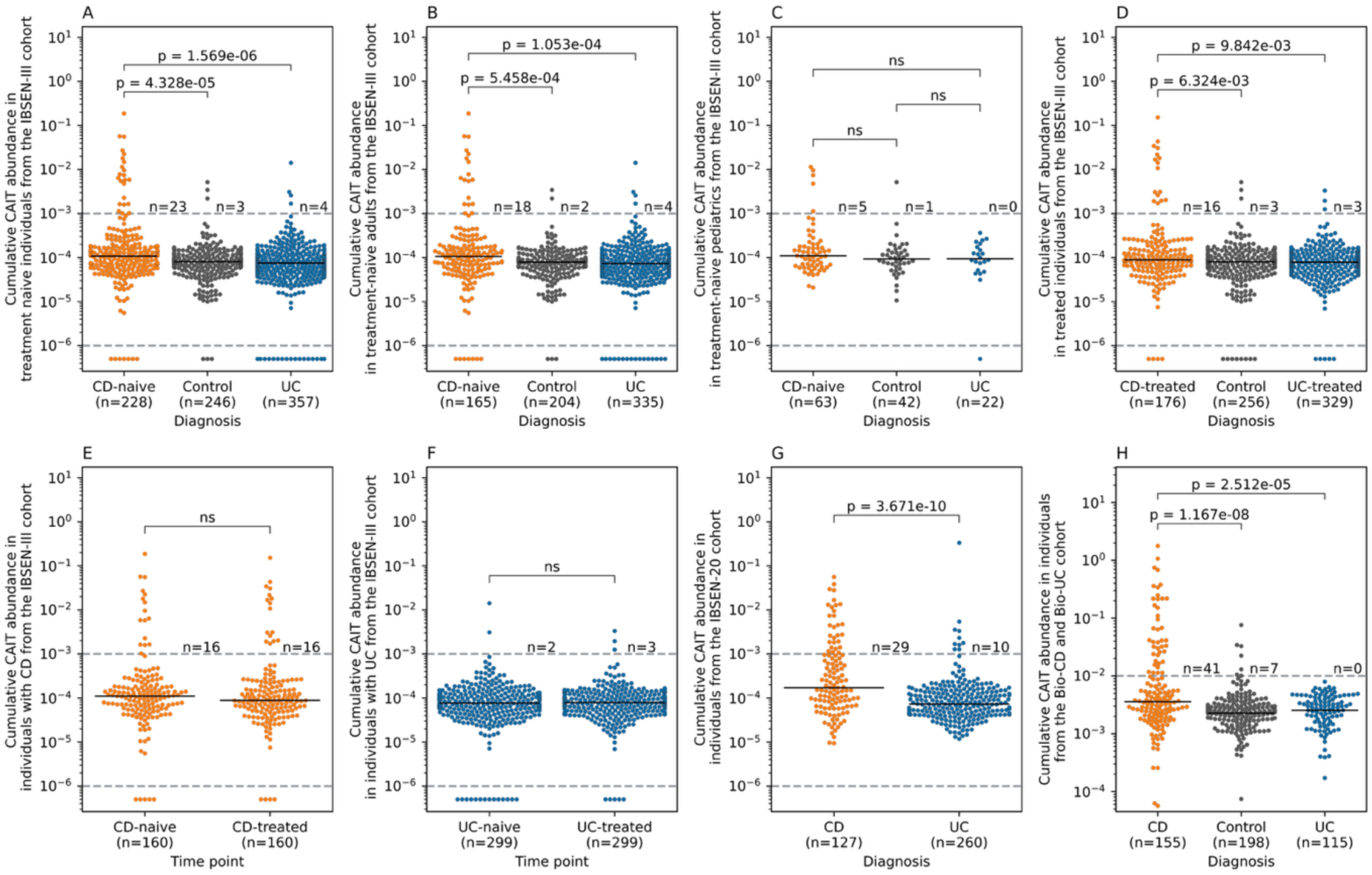
The expansion of CAIT-cells in different cohorts and phenotypic groups. (**A**) shows the expansion of CAIT cells in treatment-naive individuals with CD or UC, as well as symptomatic controls from the IBSEN-III cohort. (**B**) and (**C**) show the expansion of CAIT cells in the same phenotypic groups shown in (**A**) but separated by age, with (**B**) showing the expansion in adults (age>18 years old) and (**C**) showing the expansion in pediatric samples (age<18 years old). (**D**) illustrating the expansion of CAIT cells in treated individuals with either CD or UC, as well as symptomatic controls. (**E**) and (**F**) showing the expansion of CAIT cells in a subset of individuals from the IBSEN-III cohort with paired measurements of their T cell repertoires before and after treatment. (**G**) depicting the expansion of CAIT cells in individuals with CD or UC from the IBSEN-20 cohort. (**H**) illustrating the expansion of CAIT cells in individuals with CD relative to UC and healthy controls from the BCBC cohort. In panels (**E**) and (**F**), the expansion of CAIT cells before and after treatment was compared using the paired Wilcoxon test, while in all other panels, the two-sided Mann–Whitney U test was used to compare the expansion of CAIT cells among the different groups.

To quantify the effect of treatment on the expansion of CAIT cells, we compared their expansion in treated individuals, which recapitulated the findings observed in treatment-naive individuals (**Fig. 2D**). Indeed, by focusing on only individuals with paired measurements, *i.e.,* before and one year after treatment, we observed that CAIT cells had a comparable level of expansion in treated and treatment-naive individuals with CD (**Fig. 2E**) or UC (**Fig. 2F**). This indicates that treatment had a minor impact on the expansion of CAIT cells. Using the TRA repertoire of the IBSEN-20 cohort, we observed a significant expansion of CAIT in individuals with CD relative to individuals with UC (**Fig. 2G**). This was also replicated in the German BCBC cohort, which showed a significant expansion of CAIT cells in individuals with CD relative to healthy controls and individuals with UC (**Fig. 2H**). Thus, by profiling the T cell repertoire of three different cohorts from different geographical locations and using different methodologies, we observed a significant expansion (P_meta CD vs. controls_ = 1.4x10^-11^; P_meta CD vs. UC_ = 1.57 x 10^-17^) of CAIT cells in individuals with CD relative to UC, corroborating previous findingas^11^.

### The expansion of CAIT cells is higher in ASCA^+^ individuals with CD and individuals with ileal involvement and penetrating disease behavior

After validating the robustness of the CAIT signal across different cohorts, we aimed to investigate subphenotypes associated with a higher CAIT-expansion, focusing on individuals from the IBSEN-III cohort. Across treatment-naive and treated individuals, CAIT cells were significantly expanded in individuals with ileal involvement, *i.e.,* ileal and ileocolonic CD (**Fig. 3A** & **Fig. 3B**). This location-specific expansion was not affected by medication as the expansion was comparable in the same individuals before and after treatment (**Fig. 3C**). Disease behavior also correlated with CAIT expansion where it was higher in individuals with stricturing disease relative to individuals without a stricturing or a penetrating disease either at the treatment-naive or the treated stage (**Fig. 3D** & **Fig. 3E**). Further, the expansion of CAIT cells was comparable in individuals with CD but without a stricturing or penetrating disease, and controls. This indicates that the severity and location of the disease are major factors governing the expansion of CAIT cells and that treatment had a minor impact on the expansion of these cells (**Fig. 3F**). Besides that, ASCA status strongly correlated with the expansion of CAIT cells only in individuals with CD, which was evidenced at the IgG (**Fig. 3G**) and IgA (**Fig. 3H**) levels as well as when considering either of them (**Fig. 3I**).

**Figure 3:**
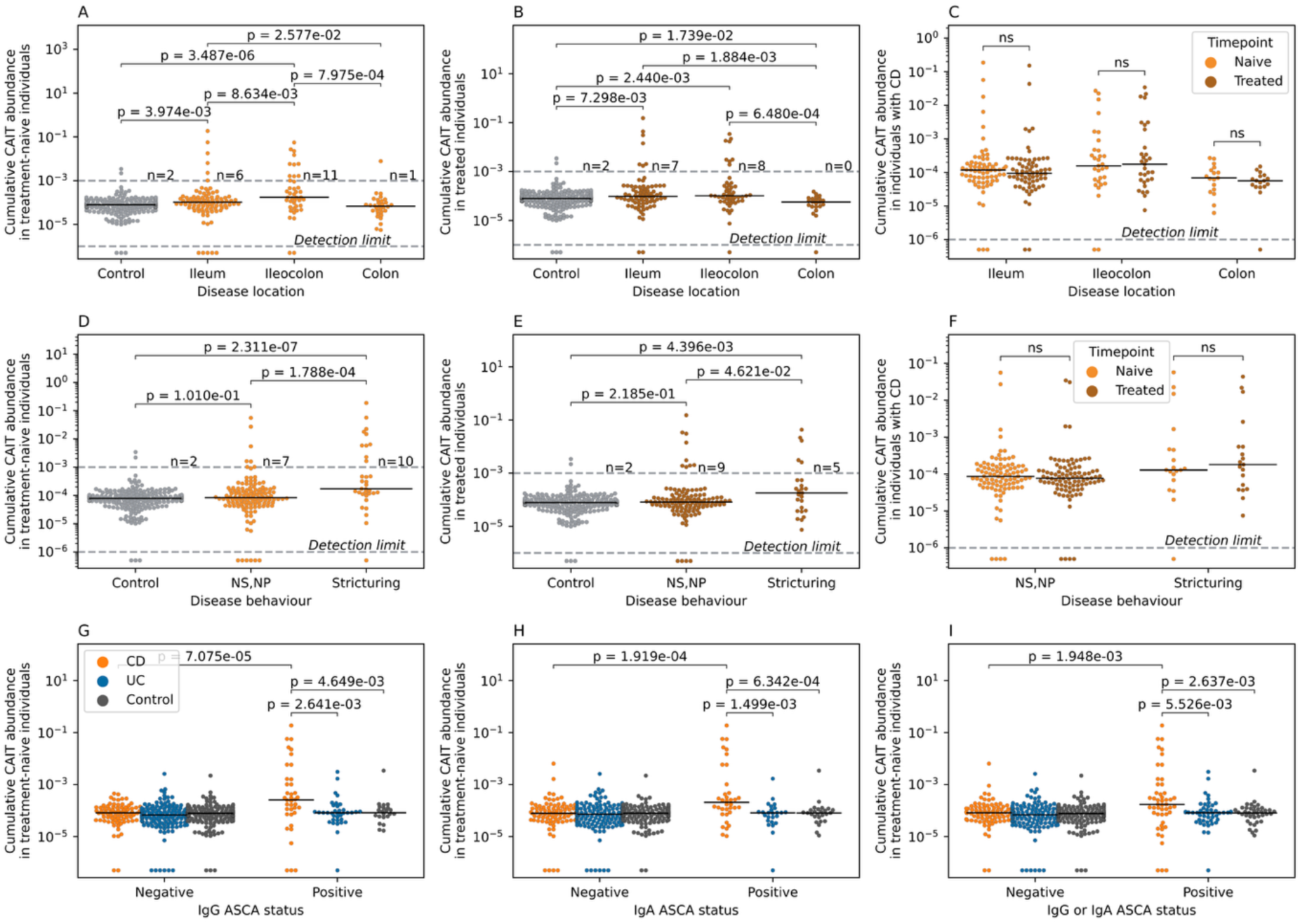
Subphenotypes and serological markers associated with high levels of CAIT expansion in individuals from the IBSEN-III cohort. (**A**) and (**B**) show the expansion of CAIT cells in treatment-naive (**A**) and treated individuals (**B**) with different forms of CD and symptomatic controls. (**C**) shows the expansion of CAIT cells in individuals with CD before and after treatment using paired measurements from the same individuals. (**D**) and (**E**) show the expansion of CAIT cells in symptomatic controls and individuals with CD who suffer from different disease behaviors. (**F**) illustrate the minor impact of treatment on the expansion of CAIT cells in individuals with different disease behaviors. (**G**) shows the expansion of CAIT cells in ASCA^+^ individuals as measured via IgG, while (**H**) depicts the same relationship but according to IgA-based measurements. Lastly, (**I**) depicts the relationship between ASCA-positivity and the expansion of CAIT cells, by defining positivity as either IgG or IgA positive.

### Mucosal-associated invariant T (MAIT) cells are significantly reduced in the blood of individuals with IBD relative to symptomatic controls

A reduction in the expansion of MAIT cells has been previously reported in individuals with IBD^11^. To investigate if this effect is related to medication-intake or the underlying disease, we compared the expansion of MAIT cells, defined as TRAV1-2+TRAJ33^+^ clonotypes, in treatment-naive and treated individuals from the IBSEN-III cohort. The expansion of MAIT cells was reduced in individuals with CD or UC but was comparable between treatment-naive and treated individuals (**Fig. S3A**). While the expansion of MAIT cells was comparable between males and females with UC or in controls, it was lower in males with CD relative to females with CD (**Fig. S3B**). Across the different diseases, the abundance of MAIT cells negatively correlated with age, indicating that age, biological-sex, and disease status can all influence the expansion of MAIT cells, and that treatment has a minor impact on the expansion of these cells.

### Hypothesis-free statistical analyses confirm previous findings and identify novel clonotypes that are associated with either CD or UC

Next, we aimed to identify other clonotypes associated with either CD or UC using a hypothesis-free statistical association framework (**Methods**). This analysis revealed 38, 72, and 13 clonotypes that were associated with CD and 35, 70, and 1 clonotypes that were associated with UC in the IBSEN-III cohort, the IBSEN-20, or the BCBC cohort, respectively. A common theme among the different sets was the detection of multiple CAIT-like clonotypes, *i.e.,* TRA chains that followed the same CAIT motif in terms of V and J gene usage and CDR3 sequence. Specifically, 2 out of the 38 CD-associated clonotypes identified from the IBSEN-III cohort, 7 out of the 72 CD-associated clonotypes identified from the IBSEN-20 cohort, and 2 out of the 13 CD-associated clonotypes identified from the BCBC cohort were CAIT clonotypes, indicating the robust association of these cells with CD. However, these multiple sets of CD-associated clonotypes did not show a robust overlap among each other (**Fig. S1A**). A similar pattern was seen among the different sets of UC-associated clonotypes (**Fig. S1B**). This could have multiple explanations, such as the stage of the disease, where different clonotypes are involved in the pathology of the disease at different stages, *e.g.,* the early stage observed in the IBSEN-III cohort relative to the late stage observed in the IBSEN-20 cohort. Alternatively, this can be attributed to differences in the sample-size among the different cohorts and hence differences in the statistical power, or a combination of these two factors.

To extend our analysis to rarer disease-associated clonotypes that we were not able to identify statistically because of the relatively small sample size of each cohort, we performed seeded clustering (**Methods**). This enabled us to identify clonotypes with a similar sequence and directionality but a lower magnitude of expansion than the clonotypes identified from the main analysis. This extended the number of CD-associated clonotypes to 111, 230, and 240 clonotypes arranged into 38, 72, and 13 meta-clonotypes derived from the IBSEN-III, the IBSEN-20, and the BCBC cohorts, respectively. Similarly, this extended the number of UC-associated clonotypes to 122, 340, and 3 clonotypes, arranged into 35, 70, and 1 meta-clonotypes, respectively. Still, limited overlap was observed between the CD-associated clonotype sets (**Fig. S2A**) as well as the UC-associated clonotype sets (**Fig. S2B**).

Before we investigated these clonotype sets further, we aimed to validate their expansion in their respective phenotypes, *e.g.,* CD-associated clonotypes in individuals with CD, among the three discovery cohorts. The expansion of CD-associated meta-clonotypes identified from the treatment-naive samples from the IBSEN-III cohort (CD_IBSEN_III) was significantly higher in individuals with CD relative to symptomatic controls and individuals with UC from the treatment-naive IBSEN-III cohort (**Fig. 4A**). Within the same cohort, the expansion of CD-associated meta-clonotypes identified from the IBSEN-20 cohort (CD_IBSEN_20) was significantly higher in individuals with CD relative to the other two groups (**Fig. 4B**). Additionally, the expansion of CD-associated meta-clonotypes identified from the BCBC cohort (CD_BCBC) was higher in individuals with CD relative to individuals with UC and controls (**Fig. 4C**). These findings indicate that the expansion of three sets of CD-associated meta-clonotypes, *i.e.* CD_IBSEN_III, CD_IBSEN_20 and CD_BCBC was significantly higher in individuals with CD included in the IBSEN-III cohort relative to individuals with UC and symptomatic controls. The same pattern was observed when comparing the expansion of these meta-clonotypes sets in the other two discovery cohorts, namely, the IBSEN-20 cohort (**Fig. 4D, 4E**, and **4F**) and the BCBC cohort (**Fig. 4F, 4G**, and **4H**). This indicates that these sets of CD-associated meta-clonotypes are robustly associated with CD and are an immunological fingerprint for CD.

**Figure 4:**
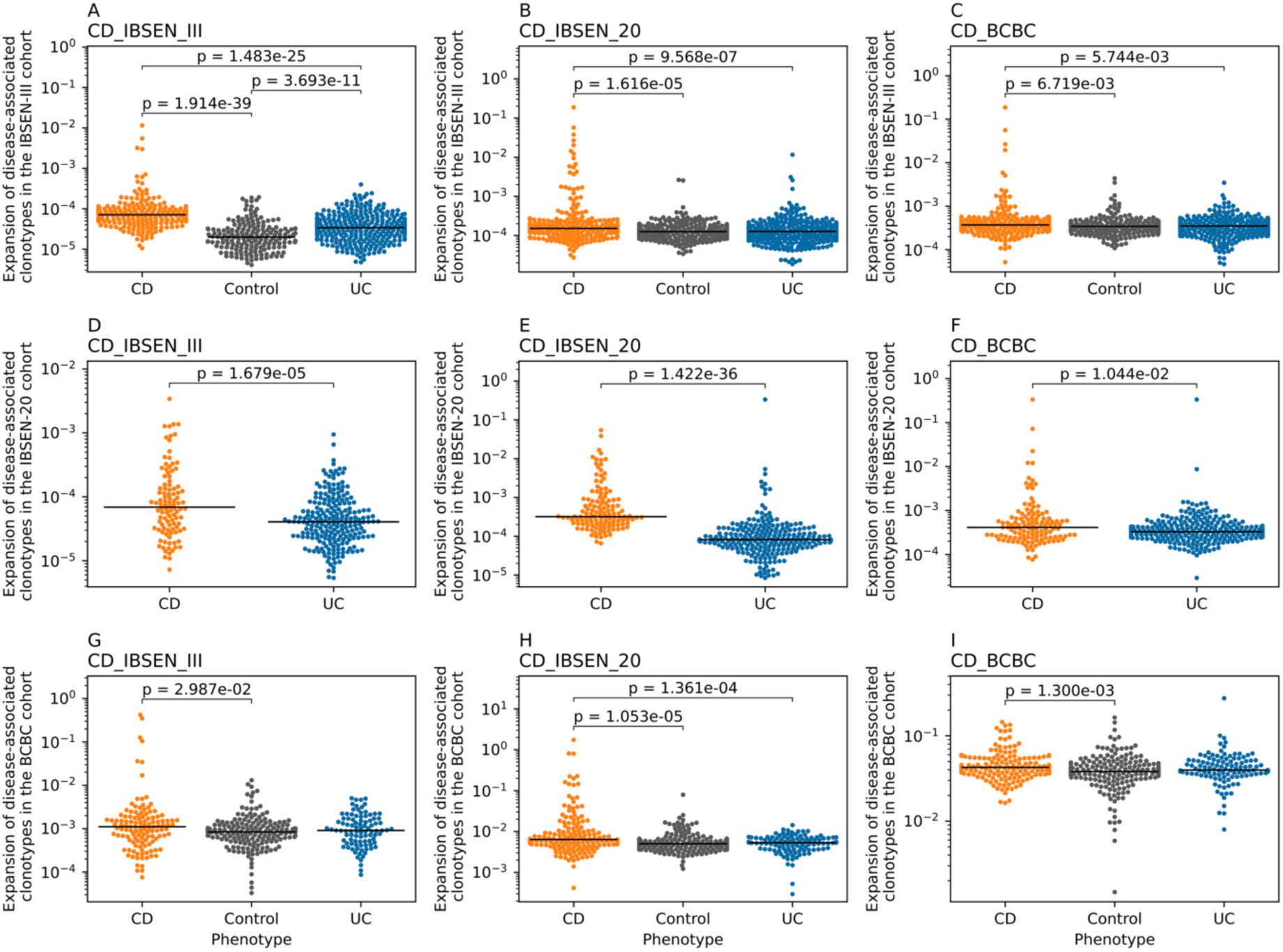
The expansion of the different CD-associated clonotypes sets in three cohorts, namely, IBSEN-III, IBSEN-20, and BCBC. (**A**), (**B**) and (**C**) show the expansion of the three CD-associated clonotype sets identified by analyzing the treatment-naive IBSEN-III cohort (CD_IBSEN_III), the IBSEN-20 cohort (CD_IBSEN_20), and the BCBC cohort (CD_BCBC cohort) in the treatment-naive IBSEN-III dataset. Similarly, (**D**), (**E**), and (**F**) show the expansion of the three CD-associated sets in the IBSEN-20 dataset, while (**G**), (**H**), and (**I**) show the expansion of these CD-associated clonotypes in the BCBC cohort.

There were notable discrepancies among the identified UC-associated meta-clonotype clonotype sets. Using the BCBC cohorts, we were able to identify only one meta-clonotype as associated with UC, potentially due to the small sample size (n=115 individuals). Hence, we focused our analysis on two UC-associated meta-clonotype sets: the first set was derived from the treatment-naive IBSEN-III cohort (UC_IBSEN_III) and the second from IBSEN-20 (UC_IBSEN_20). Within the IBSEN-III dataset, the expansion of the UC_IBSEN_III meta-clonotype set was significantly higher in individuals with UC relative to individuals with CD and symptomatic controls (**Fig. S4A**). However, the expansion of the UC_IBSEN_20 set was significantly higher in symptomatic controls than in individuals with CD or UC (**Fig. S4B**). Similarly, the expansion of the UC_IBSEN_III set was comparable in individuals with UC and CD included in the IBSEN-20 cohort (**Fig. S4C**), but the expansion of the UC_IBSEN_20 set was higher in individuals with UC relative to individuals with CD from this cohort (**Fig. S4D**). Lastly, within the BCBC cohorts, the UC_IBSEN_III meta-clonotype was predominantly expressed in individuals with UC (**Fig. S4E**), while the UC_IBSEN_20 showed the highest expansion in healthy controls relative to individuals with either CD or UC (**Fig. S4F**). Thus, within the two sets, the UC_IBSEN_III was significantly expanded in individuals with UC in its discovery cohort (IBSEN-III) and an independent validation cohort (BCBC). On the contrary, the UC_IBSEN_20 set was only expanded in individuals with UC in its discovery cohort (IBSEN-20) and neither of the two other cohorts.

This might be attributed to two interwoven reasons: first, the type of statistical comparisons used to identify UC-associated clonotypes between the IBSEN-III and the IBSEN-20. In the former, *i.e.,* the IBSEN-III cohort, UC-associated clonotypes were identified by comparing the repertoire of individuals with UC to that of controls; meanwhile, in the latter, *i.e.,* IBSEN-20, the clonotypes were identified by comparing the repertoire of individuals with UC to that of CD. Hence, the clonotypes identified from the IBSEN-20 cohort might represent a non-CD signal instead of a set of clonotypes that are associated with UC. On top of this, individuals in the IBSEN-20 cohort are generally older and have a more advanced disease course and a more complicated treatment trajectory, which might weaken or bias the disease-signal in individuals with UC.

### Meta-analysis enables the identification of a robust set of CD- and UC-associated clonotypes

After identifying and validating the expansion of the different CD-associated meta-clonotypes and a subset of the UC-associated meta-clonotypes, we aimed to integrate and unify these sets. Thus, we performed a meta-analysis across the three sets (**Methods**); specifically, for each clonotype belonging to the union of the CD- or UC-associated clonotypes, we calculated an association P-value using Fisher’s exact test in each of the three discovery cohorts. Subsequently, we combined the calculated P-values using Fisher’s combined probability test and, lastly, corrected for multiple testing using the Benjamini-Hochberg procedure (**Methods)**. Focusing on the three CD-associated clonotypes and the approach outlined above, we identified 25 clonotypes that were associated with CD (adjusted P-value <0.05). Seven out of these 25 clonotypes (∼28%) were CAIT clonotypes, corroborating previous findings about the relevance of these cells to the pathology of CD. We used the same meta-analysis-based framework to identify clonotypes that are associated with UC, focusing on the two UC-associated clonotype sets identified from the IBSEN-20 cohort and the IBSEN-III cohort, which enabled us to identify 76 public TRA clonotypes that were associated with UC. We also repeated the same process but excluded the IBSEN-20, which enabled the identification of 27 clonotypes that were associated with UC.

Before we investigated these new clonotype sets, we aimed to validate their expansion using an independent test dataset. To this end, we used a previously published dataset by Rosati *et al.*^11^, which contained the TRA repertoire of 120 individuals with CD, 47 with UC, and 100 healthy individuals. The expansion of the CD-associated clonotype set identified via the meta-analysis described above (n=25) was significantly higher in individuals with CD relative to healthy controls and individuals with UC (**Fig. 5A**). Confirming that these clonotypes capture a reproducible fraction of the immune signature of CD. We had two UC-associated clonotype sets, which were derived by including the IBSEN-20 in the analysis (n=76; set1) or by excluding IBSEN-20 from the analysis (n=27; set2). The expansion of set 1 was higher in healthy controls relative to individuals with CD and individuals with UC (**Fig. 5B**). The expansion of set 2 was significantly higher in individuals with UC relative to healthy controls (**Fig. 5C**) despite having a smaller number of clonotypes (n=27). These findings indicate that the identified clonotypes are specific to CD and UC, respectively.

**Figure 5:**
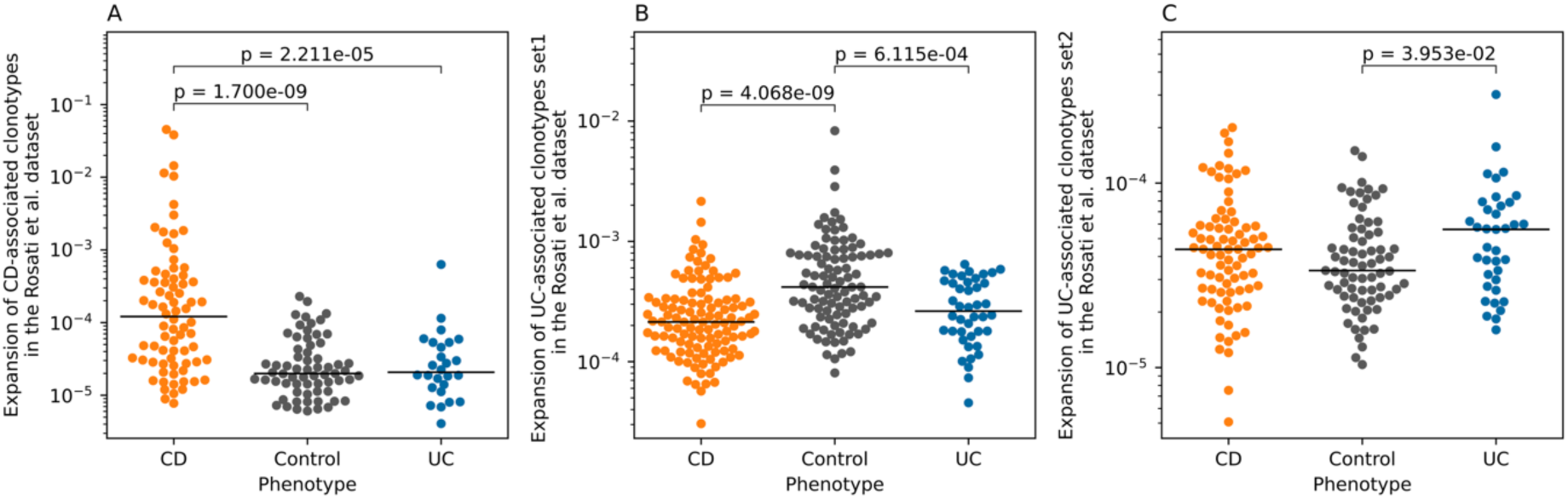
The expansion of the identified CD- and UC- associated clonotypes sets using an independent test dataset^11^. (**A**) shows the expansion of the CD-associated clonotypes in individuals with CD relative to individuals with UC and healthy controls. (**B**) and (**C**) show the expansion of the two sets of UC-associated clonotypes, *i.e.*, set 1 and set 2 described above, respectively, in individuals with either CD or UC as well as healthy controls.

### CD- and UC- associated clonotypes belong to multiple distinct clusters

After validating the identified CD- and UC- associated clonotypes using a meta-analysis as well as across independent test datasets, we aimed to understand the relationship among these clonotypes and their overlap with previously identified disease-associated clonotypes^9^. Given that we performed our meta-analysis on initial hits prior to seeded clustering, we augmented these sets with their corresponding meta-clonotypes and then performed a graph-based analysis (**Methods**). Starting with CD-associated meta-clonotypes, we observed multiple distinct clusters (**Fig. 6A**), the largest of them belonging to CAIT cells as they share the same V and J gene combination and CDR3 amino acid motif (**Fig. 6B**). The second biggest cluster has a *TRAV29-01* and *TRAJ06-01* based combination and the following CDR3 amino acid motif (CAASA**GGSYIPTF) (**Fig. 6C**). Lastly, the third biggest cluster showed a MAIT-like VJ-recombination that is derived from *TRAV01-02* and *TRAJ33-01* as well as a conserved CDR3-amino acid motif that only varied in a single amino acid position (**Fig. 6D**).

**Figure 6:**
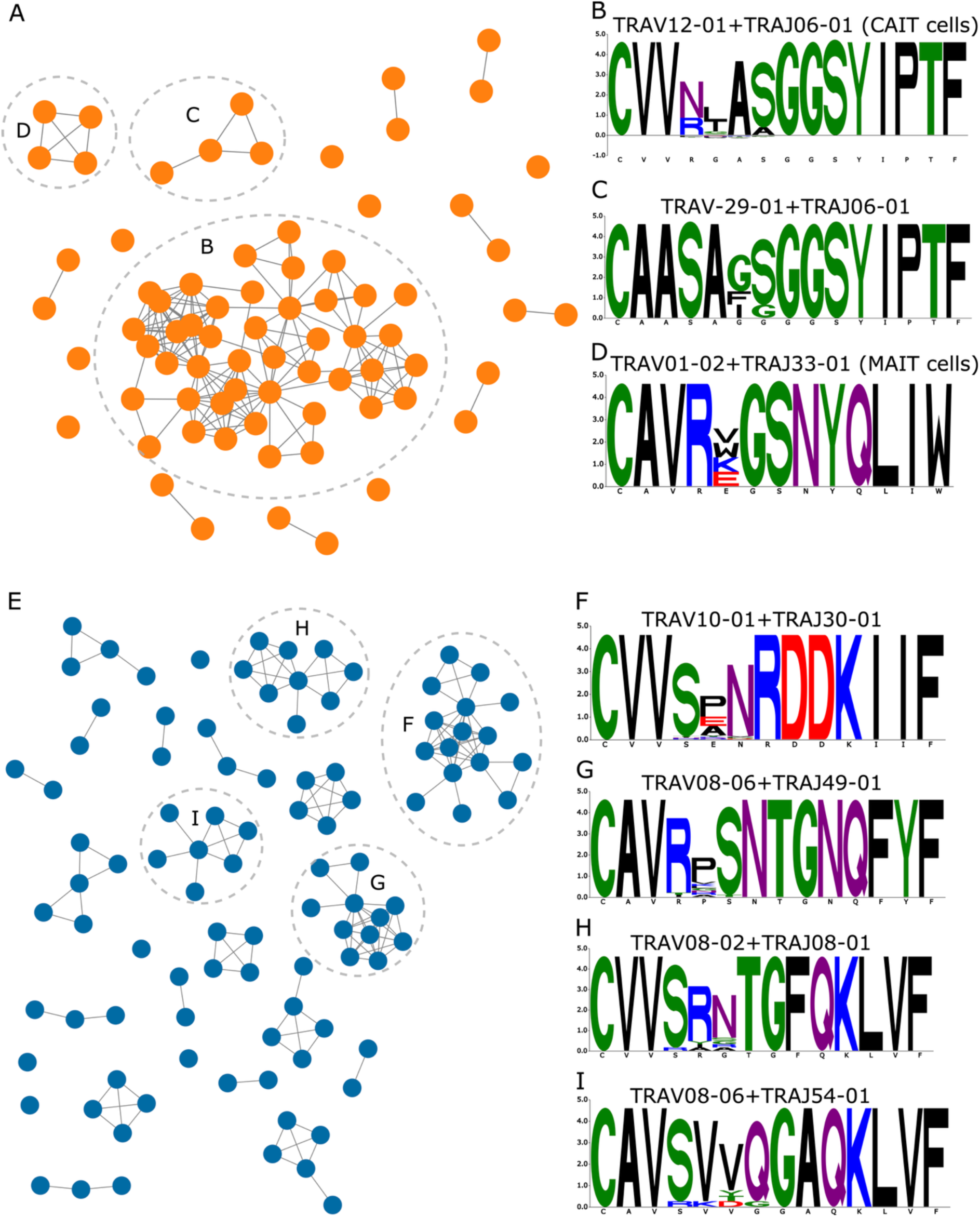
A network analysis of CD- and UC-associated clonotypes (set 2) identified from the meta-analysis conducted across the IBSEN-III, the IBSEN-20, and the BCBC cohort. (**A**) it is a graph-based representation of CD-associated clonotypes where nodes represent clonotypes, while edges represent similarity among these clonotypes; specifically, two nodes are connected if they have the same V and J genes and their CDR3 is different by only 1 Hamming distance. (**B**-**D**), it is a motif representation of the CDR3 amino acid sequence of the three biggest CD-associated clonotypes depicted in (**A**). (**B**) it is a network representation of the UC-associated clonotypes (set 2) identified from the cross-cohorts meta-analysis. (**F**-**I)** depict the CDR3-amino acid motif for the four biggest UC-associated clusters.

There were also multiple distinct clusters identified from the UC-associated meta-clonotypes (**Fig. 6E**). The biggest of these clusters were derived from a combination between *TRAV10-01* and *TRAJ30-01* with a CDR3-motif that predominantly differed in only one amino acid position (**Fig. 6F**). There were also multiple clusters derived from the TRAV08 family, for example, the second biggest cluster was derived from a combination between *TRAV08-06* and *TRAJ49-01* and CDR3-amino acid motif that differed in one amino acid position (**Fig. 6G**). The third biggest cluster was a composite of a VJ recombination between the *TRAV08-02* and *TRAJ08-01* segments and a CDR3-amino acid motif that differed on only two amino acid positions (**Fig. 6H**). The last cluster with more than seven clonotypes was derived from a *TRAV08-06* and *TRAJ54-01* based recombination; the members of this cluster showed a conserved CDR3 motif with a potential variation in one to two positions (**Fig. 6I**).

After identifying these clusters, which indicate a focused immune response toward multiple distinct antigens, we aimed to investigate the antigenic specificity of these clonotypes. Using yeast-specific TCR sequences datasets^9^, we detected multiple overlaps with the set of CD-associated clonotypes, particularly with CAIT cells (n=14 clonotypes). There was no overlap with UC-associated clonotypes, which is consistent with the fundamental role of anti-yeast responses in CD but not in UC. Through the utilization of T cell Receptor-Antigen InTeractions databases (TRAIT)^23^, which is a recently published dataset of TCR sequences with their antigenic specificity, we were able to identify the antigenic specificity of one CD-associated clonotype that was restricted to a SARS-CoV-2 peptide presented by the HLA-A*02 protein. Several factors can explain this, such as cross-reactivity between the antigen(s) recognized by these TRA clonotypes and SARS-CoV-2, noise in the public annotation database, among others. Similarly, we were not able to hypothesize on the antigenic specificity of any of the UC-associated clonotypes.

## Discussion

Several studies previously aimed at identifying antigens and risk factors potentially causing IBD^9,10,24^, although some risk factors have been identified, *e.g.,* antibiotic intake^25^, and infectious mononucleosis^26^, the etiology of the disease remains far from being understood. While TCR-Seq does not enable the direct identification of these antigens, it can pinpoint their trace on the adaptive immune system by identifying clonotypes recognizing these antigens^14,19^. Focusing on the TRA repertoire, which is less understood and studied than the TRB repertoire, the discovery of CAIT cells by Rosati *et al.*^11^ using hundreds of TRA samples represented a first step toward identifying specific TRA clonotypes associated with CD. This highlighted the need for profiling the T-cell repertoire of large cohorts to identify these disease-associated exposure markers. It should be stressed that CAIT cells can currently only be profiled using TRA and not TRB because of their semi-invariant TRA chains, with their TRB repertoire remaining largely under-investigated.

Here, we aimed not only to increase the sample size but also to include multiple cohorts spanning different stages of the disease. This not only enabled us to perform within-cohort analyses but also to perform a first-of-its-kind meta-analysis across these different cohorts. To this end, we were able to identify clonotypes that were specific to IBD in its early stage, late stage, and to the disease across all stages as revealed by the meta-analysis. One of the consistent signals across the different stages was the known CAIT signal. This illustrates that these cells are a stable, robust marker of CD across the entire disease trajectory, particularly in individuals with ileal and ileocolonic CD. Further, it also highlights that the expansion of CAIT cells is not impacted by the different therapeutic trajectories and surgeries. Hence, deconvolving the antigenic specificity of these cells and discovering their role in the disease is a promising strategy to understand the etiology of CD.

Large-scale TRB repertoire profiling studies that span thousands of individuals have identified hundreds to thousands of TRB clonotypes that are associated with the disease and simultaneously were able to discover their HLA restriction^27^. While our study represents the largest TRA analysis study published-to-date, its sample size is smaller than these large TRB-based studies, and hence, its power to identify disease-associated clonotypes is limited. One of the reasons that enabled us to identify CAIT with our relatively small sample-size is their nature as unconventional T cells coupled with a large effect-size. These T cells are not restricted to HLA proteins, which are highly polymorphic, but to the CD1d protein, which is monomorphic; hence, a relatively small sample size was required to associate them with the disease.

As large-scale bulk TCR repertoire profiling becomes the default method to identify disease-associated clonotypes^17,19,28–30^, the more urgent it becomes to identify the phenotypes, functions, and pathological roles of the identified disease-associated clonotypes. Different research directions can be followed to discover and investigate these different aspects, such as the development of animal models to study the therapeutic potential of the targeted depletion of disease-associated clonotypes^5,31,32^. Additionally, bulk TCR-Seq identifies a particular chain, *e.g.,* TRA or TRB, that is associated with these diseases instead of paired αβ T cell receptors. Therefore, generating paired TCRs would be a prerequisite step for disentangling the antigenic specificity of these clonotypes. Several methods can be used to generate the pairing, such as pairSEQ^33^ and TIRTL-Seq^34^, as well as single-cell RNA sequencing^11^. The combination of these methods will be an important next step, as pairing information is necessary to further refine clonotypes and for functional follow-up studies.

## Author contributions

H.E. and A.F. conceived and designed the study. A.K.H.M. performed data curation. A.K.H.M., E.E.K., and V.K. prepared the TRA NGS libraries from the IBSEN-III cohort. C.O., G.P., M.B.B., P.R., S.A., T.E.D., V.A.K., J.R.H., and M.L.H. collected the biomaterial from the IBSEN-III and/or the IBSEN 20 cohorts, in addition to collecting, storing, and processing clinical and metadata. M.P., D.M., S.S., B.H., and H.R. collected and profiled the TCR alpha repertoire of the BCBC cohort. H.E. and A.K.H.M. made the figures, performed the analyses, and wrote the first draft of the manuscript with input from all co-authors. All authors read and approved the final version of the manuscript.

## Data availability

Due to GDPR and consent restrictions, the datasets reported in the current study, namely, the TCR-Seq datasets of the BCBC cohort, can be obtained by submitting a project application to the PopGen 2.0 Network (https://portal.popgen.de/ ). This dataset will be kept and made available for at least 10 years from the date of publication, and the processing time of applications is approximately 6-8 weeks. Regarding the IBSEN-III and the IBSEN-20 datasets, institutional data privacy regulations prohibit the deposition of individual-level data in public repositories. Participant written consent also does not cover public sharing of data for use for unknown purposes. Upon contact with Marte Lie Høivik, an institutional data transfer agreement can be established and data shared if the aims of data use are covered by ethical approval and patient consent. The procedure will involve an update to the ethical approval as well as a review by legal departments at both institutions, and the process will typically take one to two months from initial contact. The analytical code and software used in the current study are based on publicly available tools and custom scripts developed in Python, as described in the Materials and Methods section.

## Acknowledgment

We would like to thank Sören Franzenburg, Janina Fuß, Rebekka Kraemer, Nicole Braun, Maria Eloina Figuera Basso, Anja Tanck, Xiaoli Yi, Tanja Wesse, Yewgenia Dolshanskaya, Melanie Vollstedt, and Cathrin John-Klaua from the Institute of Clinical Molecular Biology (Kiel University and University Hospital Schleswig-Holstein, Kiel, Germany) for their help with profiling the TRA repertoire of the IBSEN-III and the IBSEN-20 cohort. We would also like to thank Tanja Weise and Michael Wittig for their genotyping of the IBSEN-III and IBSEN-20 cohorts. Lastly, we would like to thank the IBSEN III study group for their support in conducting the IBSEN III study.

