## Supplementary figures for "Multi-centered T cell repertoire profiling identifies novel alterations in the immune repertoire of individuals with inflammatory bowel disease and validates previous findings"

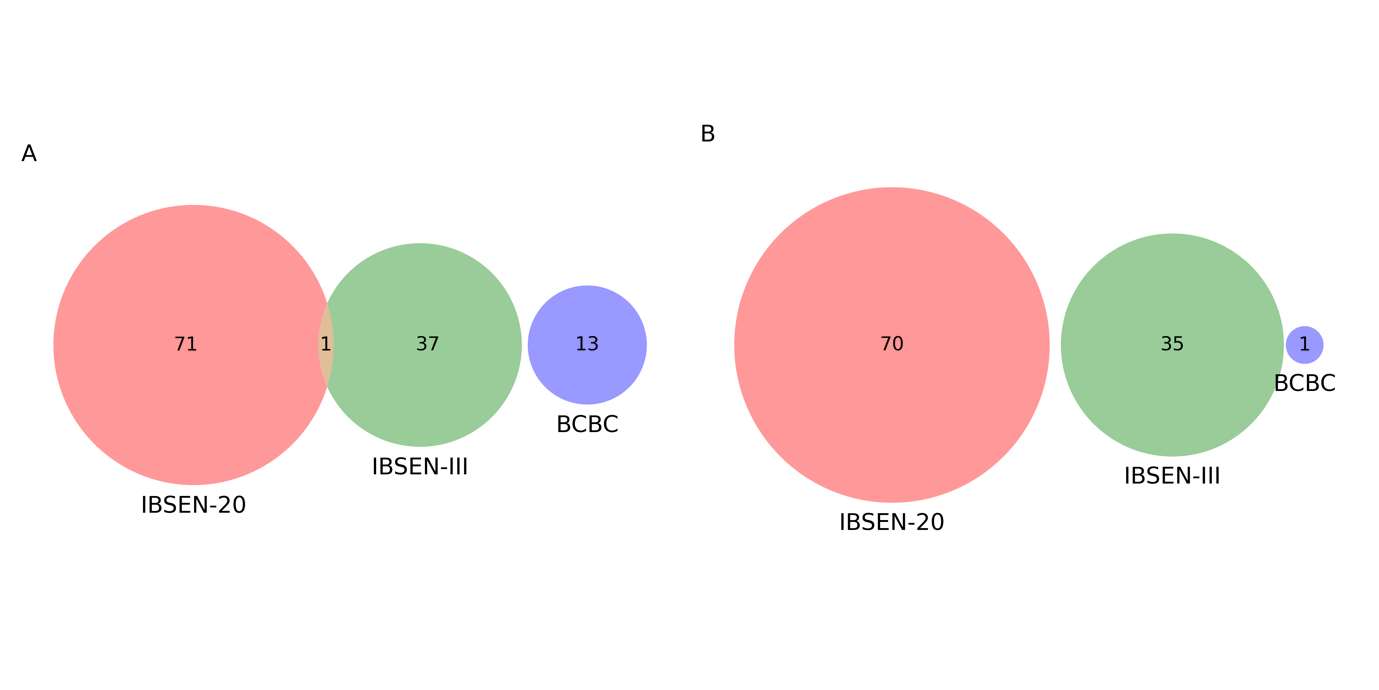

**Figure S1**: The overlap among disease-associated clonotypes identified by analyzing each cohort independently. (**A**) depicts the overlap among CD-associated clonotypes, while (**B**) depicts the overlap among UC-associated clonotypes.

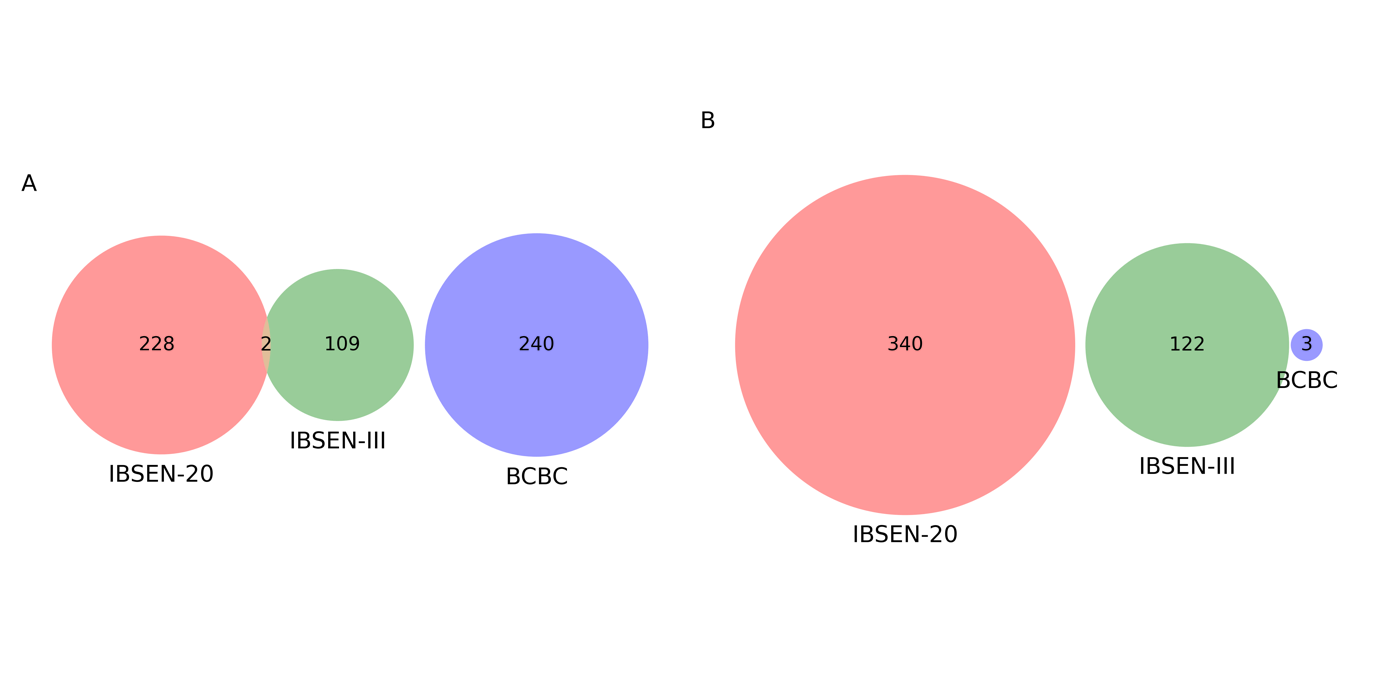

**Figure S2**: The overlap in CD- and UC- associated meta-clonotypes identified using seeded-clustering. (**A**) illustrate the overlap at the level of CD-associated meta-clonotype sets, while (**B**) shows the overlap among the UC-associated meta-clonotype sets.

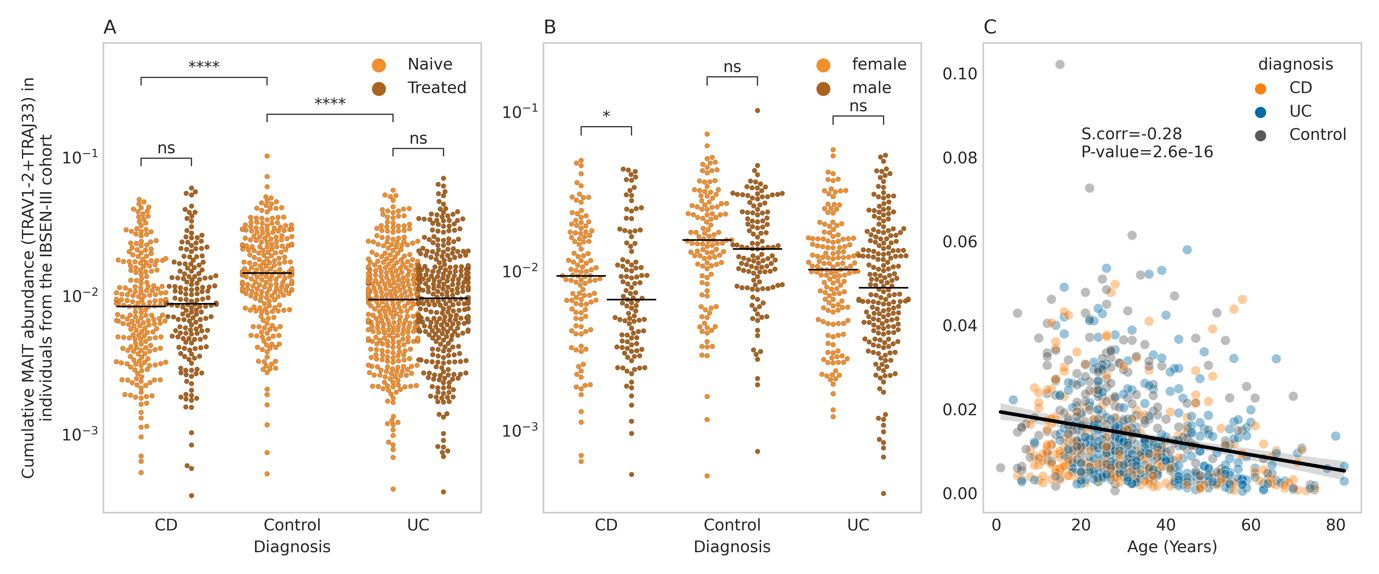

**Figure S3:** The impact of IBD, therapy, biological sex, and age on the expansion of MAIT cells. (**A**) the expansion of MAIT cells in individuals from the IBSEN-III cohorts, namely, treatment-naïve and treated individuals with CD and UC, and symptomatic controls. (**B**) shows the impact of biological sex on the expansion of MAIT cells in treatment-naïve individuals, while (**C**) shows the strong negative correlation between the expansion of MAIT cells and age in years, regardless of disease status.

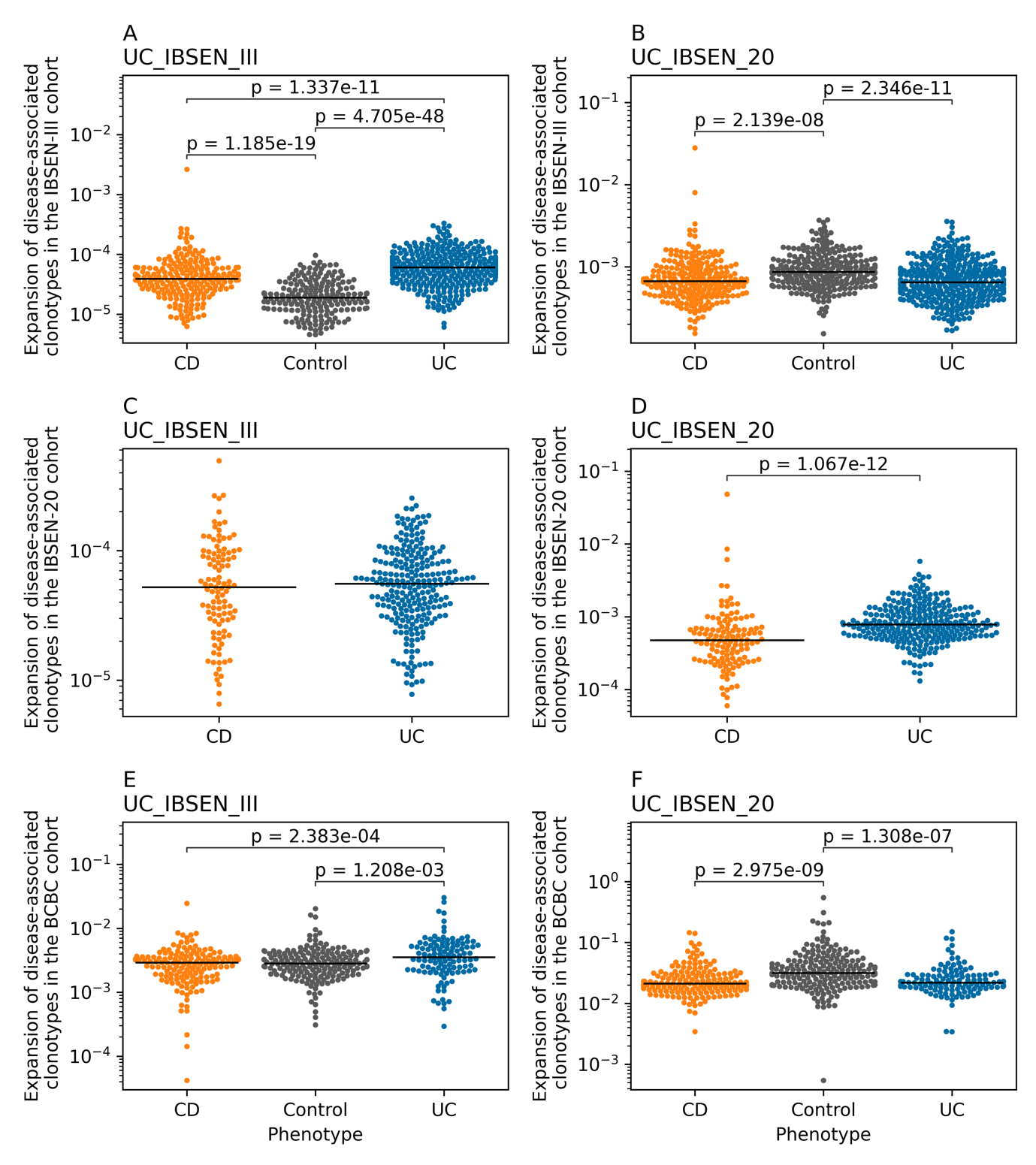

**Figure S4**: The expansion of the different UC-associated clonotypes in the three discovery cohorts, namely, IBSEN-III, IBSEN-20, and BCBC. (**A**) and (**B**) show the expansion of the UC-associated clonotype sets identified from the analysis of the IBSEN-III cohort (UC_IBSEN_III) and IBSEN-20 cohort (UC_IBSEN_20) in the TRA repertoire of IBSEN-III. (**C**) and (**D**) show the expansion of the identified UC-associated clonotype sets in the repertoire of IBSEN-20 individuals. Lastly, (**E**) and (**F**) depict the expansion of UC-associated clonotypes sets across individuals included in the BCBC cohort as well as in the included healthy controls. Across all panels and unless stated otherwise, all statistical comparisons were conducted using the two-sided Mann–Whitney U test.

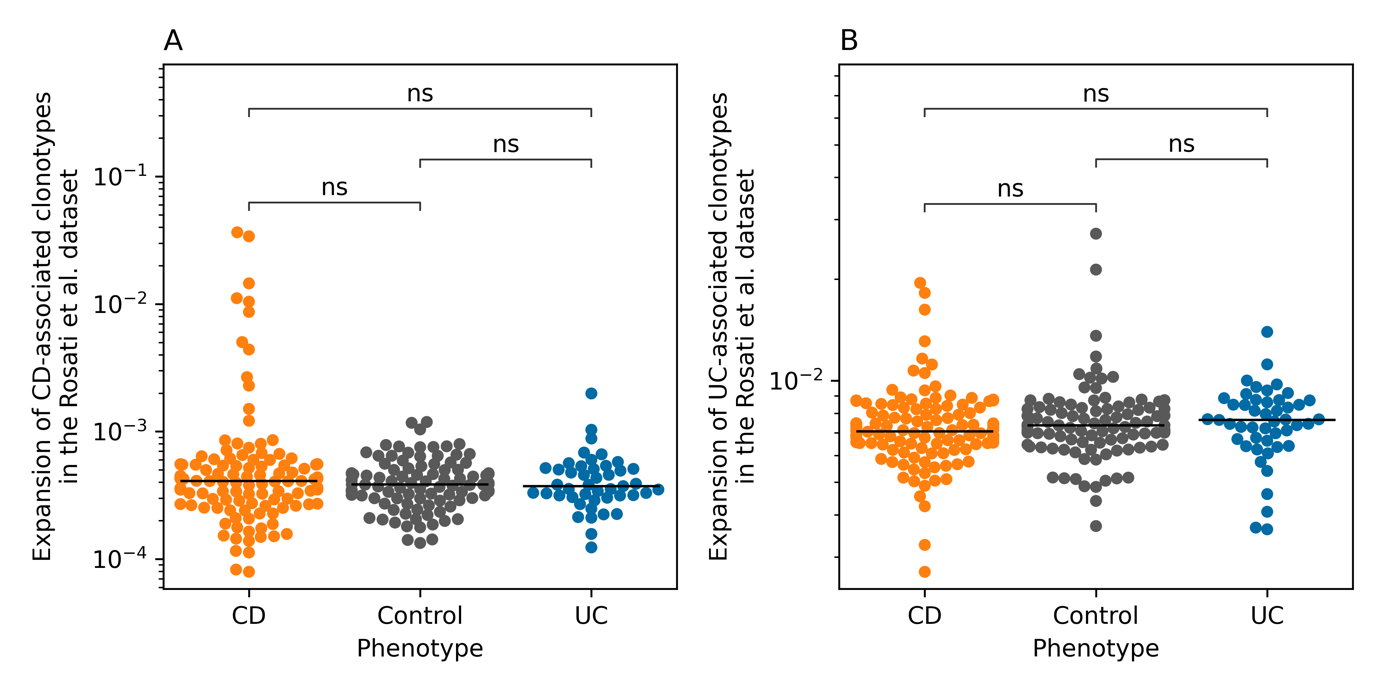

**Figure S5**: The expansion of the CD- and UC- associated clonotypes identified by integrating the TRA of the three cohorts in the Rosati et al.^11^ datasets. (**A**) shows the expansion of CD-associated clonotypes, while (**B**) shows the expansion of UC-associated clonotypes.

**Supplementary tables**

**Table S1:** Phenotypic properties of the IBSEN-III samples included in the study after removing samples with less than 1,000 functional, i.e., productive, clonotypes which are clonotypes that encode for functional TRA or TRB chains and do not contain a stop-codon or a frameshift mutation.

|  | Crohn’s disease | Ulcerative Colitis | Symptomatic controls |
| --- | --- | --- | --- |
| Total number of samples | 237 | 384 | 240 |
| Number of samples at the treatment-naïve state | 215 | 343 | 240 |
| Number of samples at the treated state | 170 | 321 | 0 |
| Number of samples with paired measurements | 148 | 280 | 0 |
| Percentage of females | 53.16% (n = 126) | 44.27% (n = 170) | 49.17% (n = 118) |
| Age at diagnosis | 32 ± 17.44 years | 36.38 ± 14.76 years | 29.10 ± 13.41 years |
| Number of pediatric cases | 64 | 23 | 41 |
| Disease location and extension at the treatment-naïve status | L1: Ileal (n= 83; 38.6%) | E1: Ulcerative proctitis (n=133; 38.77%) | **-** |
|  | L2: Colon (n=29; 13.48%) | E2: Left-sided colitis (n=67; 19.53%) | **-** |
|  | L3: Ileocolonic (n=40; 18.60%) | E3: Pancolitis (n=116;33.81%) | **-** |
|  | Unknown (n=63; 29.30%) | Unknown (n=27;7.87%) | **-** |
| Disease modifiers at the treatment-naïve status | B1: Non-stricturing, non-penetrating (n=152; 70.69%) | **-** | **-** |
|  | B2: Stricturing (n=30; 13.95%) | **-** | **-** |
|  | B3: Penetrating (n=5; 2.3%) | **-** | **-** |

**Table S2:** Phenotypic properties of the IBSEN-20 samples included in the study after removing samples with less than 1,000 functional, i.e., productive, clonotypes.

|  | Crohn’s disease | Ulcerative Colitis |
| --- | --- | --- |
| Total number of samples | 127 | 260 |
| Percentage of females | 48% (n = 61) | 50.38% (n = 131) |
| Age at diagnosis | 31.76 ± 13.73 years | 35.26 ± 11.35 years |
| Disease location and extension at the treatment-naïve status | L1: Ileal (n= 19; 14.96%) | E1: Ulcerative proctitis (n=53; 20.38%) |
|  | L2: Colon (n=36; 28.34%) | E2: Left-sided colitis (n= 92; 35.38%) |
|  | L3: Ileocolonic (n=72; 56.69%) | E3: Pancolitis (n=115; 44.23%) |
| Disease modifiers at the treatment-naïve status | B1: Non-stricturing, non-penetrating (n=49; 38.58%) | **-** |
|  | B2: Stricturing (n=41; 32.28%) | **-** |
|  | B3: Penetrating (n= 37; 29.92%) | **-** |

**Table S3:** Phenotypic properties of the BCBC cohort included in the current study as well as the matching healthy controls.

|  | Crohn’s disease | Ulcerative Colitis | Healthy controls |
| --- | --- | --- | --- |
| Total number of samples | 155 | 115 | 198 |
| Percentage of females | 58.06% (n = 90) | 38.26% (n = 44) | 52.52% (n = 104) |
| Age at sample collection | 38.24±13.41 years | 39.4±14.33 years | 44.69 ± 13.522 years |
| Disease location and extension at the treatment-naïve status | L1: Ileal  (n= 55; 35.48%) | NA | **-** |
|  | L2: Colon  (n=38; 24.51%) | NA | **-** |
|  | L3: Ileocolonic (n=53; 18.60%) | NA | **-** |
| Disease modifiers at the treatment-naïve status | NA | **-** | **-** |
|  | NA | **-** | **-** |
|  | NA | **-** | **-** |
